# Real-time demultiplexing Nanopore barcoded sequencing data with npBarcode

**DOI:** 10.1101/134155

**Authors:** Son Hoang Nguyen, Tania Duarte, Lachlan J. M. Coin, Minh Duc Cao

**Affiliations:** Institute for Molecular Bioscience, The University of Queensland, Brisbane, St Lucia, QLD 4072, Australia

## Abstract

**Motivation:** The recently introduced barcoding protocol to Oxford Nanopore sequencing has increased the versatility of the technology. Several bioinformatic tools have been developed to demultiplex the barcoded reads, but none of them support the streaming analysis. This limits the use of pooled sequencing in real-time applications, which is one of the main advantages of the technology.

**Results:** We introduced npBarcode, an open source and cross platform tool for barcode demultiplex in streaming fashion. npBarcode can be seamlessly integrated into a streaming analysis pipeline. The tool also provides a friendly graphical user interface through npReader, allowing the real-time visual monitoring of the sequencing progress of barcoded samples. We show that npBarcode achieves comparable accuracies to the other alternatives.

**Availability:** npBarcode is bundled in Japsa - a Java tools kit for genome analysis, and is freely available at https://github.com/hsnguyen/npBarcode.

## 1. Introduction

Introduced only in 2014, Oxford Nanopore Technologies (ONT) sequencing has already become an established technology for its portability and its potential for high yield data generation. In particular, it offers the ability of real-time sequencing where practitioners can analyse data streaming directly from the sequencing device, and can terminate a sequencing run once the satisfactory results are obtained. Recently, ONT introduced barcode protocols to allow pooling and sequencing multiple libraries on the sample flow cell, which further enhances the versatility of the technology. The underlying mechanism is to ligate a unique oligonucleotide sequence, or *barcode*, to the fragments of each DNA sample. Multiple samples can then be pooled together and sequenced in one flow cell. The sequenced reads can then be demultiplexed into bins by examining the barcode portions on the reads.

Several outstanding tools for demultiplexing Nanopore barcoded sequences such as poreFUME [6], Porechop [7] and Metrichor built-in demultiplexer have been developed. Of these tools, only the latter supports real-time analysis of a sequencing run, but it is only available as a cloud service. This limits the use of this technology in time-critical applications or when the users wish to perform sequencing only until sufficient data are obtained.

Here, we present npBarcode, a sensitive tool for demultiplexing barcoded MinION sequencing data in real-time. npBarcode provides the traditional command line interface and a graphical user interface. The command line interface offers a flexible environment to be integrated in with other real-time downstream analyses, *e.g.* real-time finishing genome sequence (npScarf[4]) and real-time species identification (npAnalysis[3]). The demultiplexer is also integrated into npReader’s[2], our previously developed platform for real-time analysis and visualisation of nanopore sequencing. From this mode, beside the utilities provided by npReader, one can visually monitor on the sequencing progress of each barcoded sample.

## 2. Methods

npBarcode relies on local pairwise alignment to detect the existence of a barcode sequence in each nanopore read. We used the Smith-Waterman algorithm with Gotoh improvement [5] for the alignment. Basically, the aligner attempts to align the barcode sequences within a window on both ends of a nanopore read. The read is assigned to the group of the barcode with the highest alignment score, provided that the score is greater than a threshold, and is greater than the second best alignment score by a safety distance.

npBarcode is implemented within the Japsa toolkit as a program named jsa.np.barcode. The program takes the base-called data as a stream input and demultiplexes the data into different channels containing reads that belong to the same bin. These output streams can be piped to other downstream real-time analyses. This design allows practitioners to integrate the tool into a streaming analysis pipeline of interests.

We barcoded and sequenced 8 different bacterial strains on a single MinION R9.4 flow cell using the Oxford Nanopore Technology’s 2D native barcoding kit (SQK-LSK208 + EXP-NDB002). The pool consisted of one gram-positive isolate (*Staphylococcus aureus*) and seven gram-negative isolates (four *Klebsiella pneumoniae*, one *Klebsiella quasipneumoniae*, one *Pseudomonas aeruginosa* and one *Acinetobacter baumannii*). In order to establish a ground truth benchmark for comparison of different de-multiplexing tools, we aligned the sequence reads of the eight samples to their respective assemblies to identify the original source of the reads. Due to the high level of similarity among the gram negative isolates, we could only obtain the confident assignment of reads to the *S. aureus*, a gram positive strain, versus the other gram negative strains. Hence, we set up the comparison framework as the accuracy of the tool detecting the barcode associated with the *S. aureus* sample. Figure 1 presents the sensitivity and specificity of npBarcode with differing parameters. We also obtained the results from Metrichor, porechop and poreFume with their default parameter settings. We found that npBarcode performed comparably with Metrichor and was marginally more accurate than the competitive porechop and poreFUME.

**Figure 1:**
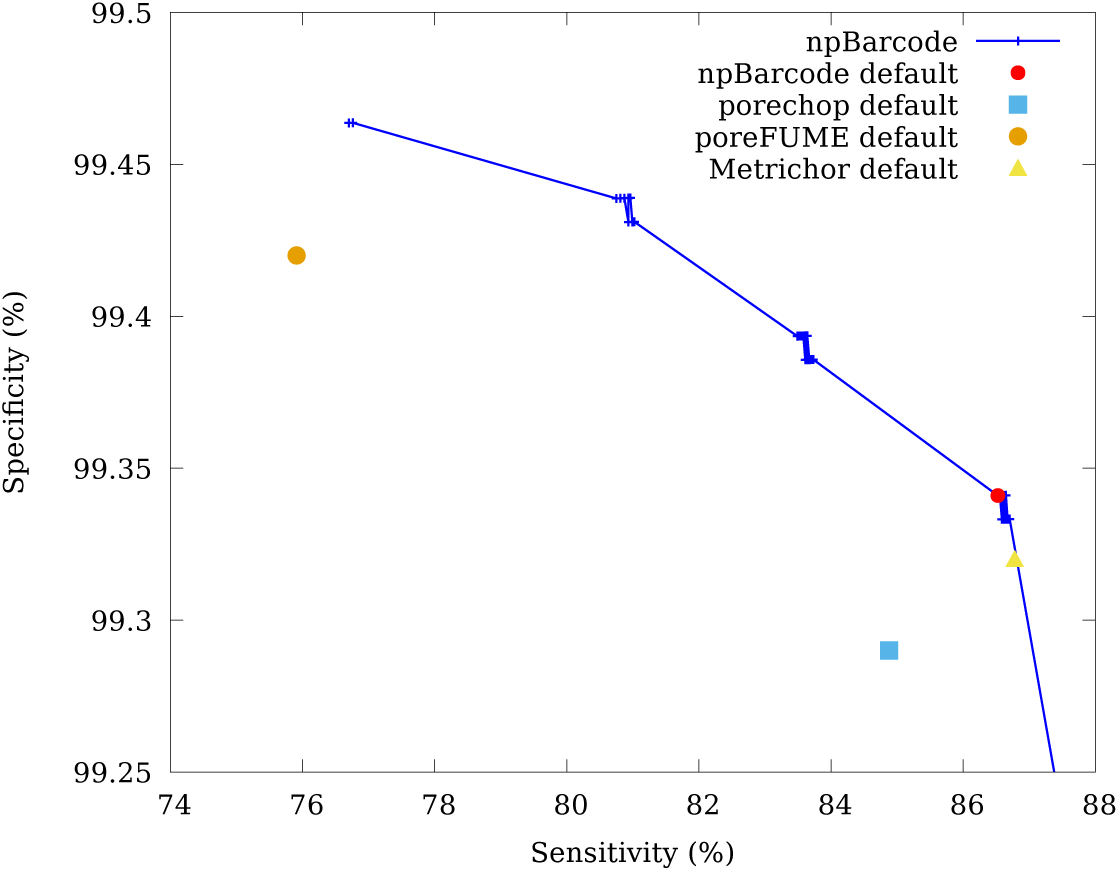
Plot of sensitivity versus specificity of npBarcode compared with existing tools.

As part of the experiment, we also integrated npBarcode into a streaming analysis pipeline where we scaffolded genome assemblies in real-time. Prior to the MinION sequencing run, we sequenced the seven gram negative samples with Illumina technology and used SPAdes [1] to assemble them into assemblies, which each had exceeding 100 contigs. The *S. aureus* sample was used as a control sample during demultiplexing, and was not used for scaffolding. As soon as a sequence read was generated, it was base-called, demultiplexed and streamed into the appropriate instance of npScarf [4] for scaffolding. This pipeline allowed us to simultaneously scaffold all seven bacterial samples while the sequencing was still in progress (Figure 2). We were able to complete the genome of one *K. pneumoniae* isolate after 16 hours of sequencing and with about 80Mb of nanopore sequencing data. While the genomes of the other six isolates were not completed due to their low proportions in the pooled sample, and the less than ideal yield of the run, they were significantly improved over time with nanopore long reads.

**Figure 2:**
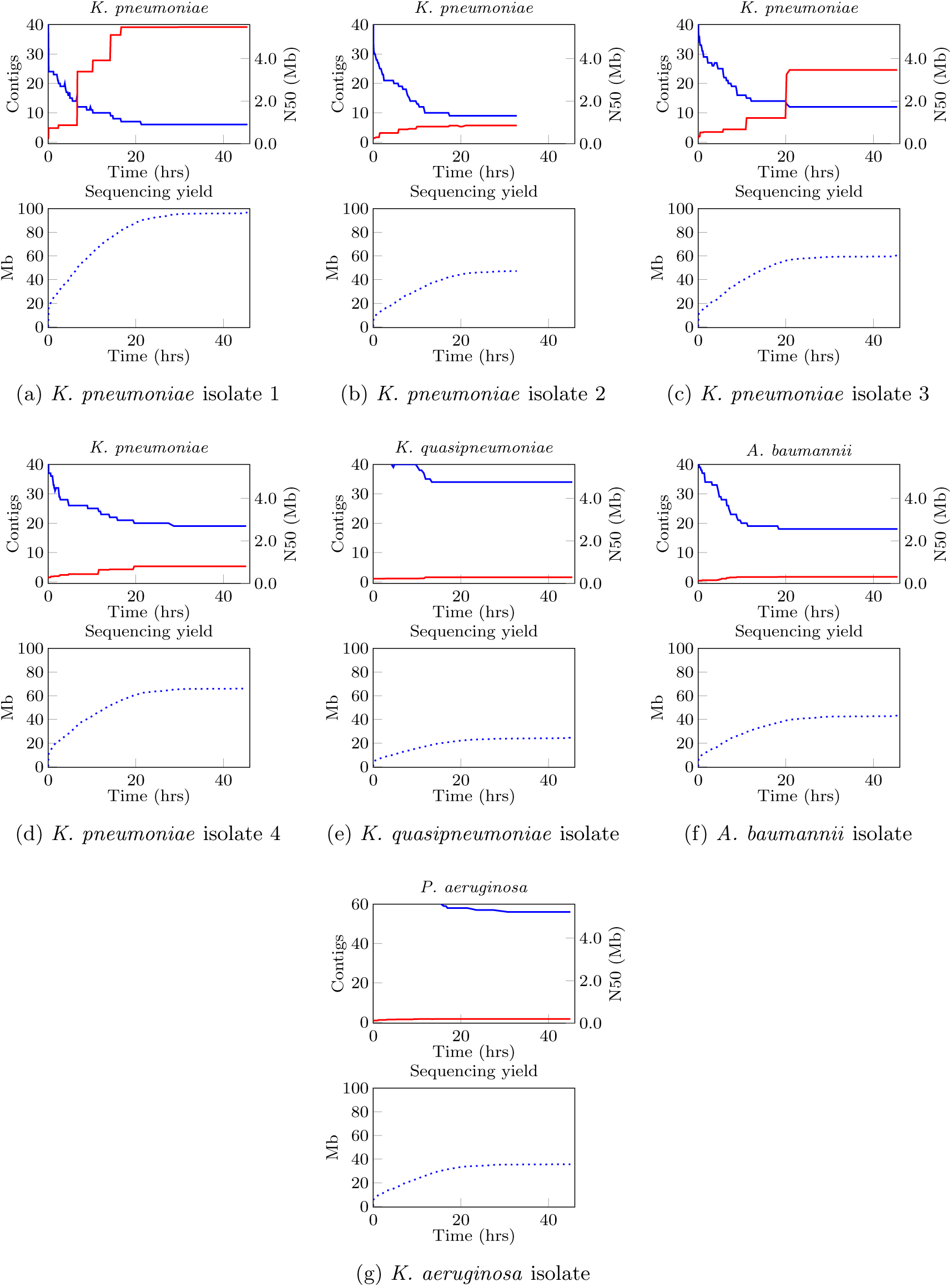
Result for running a combining pipeline of npReader, npBarcode and npScarf with ONT Native Barcoding Kit in order to scaffold genomes of 7 gram-negative samples in real-time. For each sample, the upper plot shows number of contigs (blue) and N50 (red) while the lower graph presents data yield (bases) for that particular demultiplexed sample over time.

npBarcode is also integrated into npReader’s graphical user interface. With this, users can monitor the amount of sequencing data for each pooled sample in real-time. When required, npReader provides a view showing read-count per bin in a real-time fashion. This view provides two plots that can depict the progress in an over-time and in-time manner, respectively. The first scenario would offer more general progress viewing by showing the read count of the whole process while the lower graph figures the exact number of binned reads at a particular time point. An example of the visualisation is shown in Figure 3. Overall it reflects the demultiplex functioning appropriately with the majority of reads falling in the bins corresponding to the samples used for the barcode sequencing. Only an negligible amount of reads are mis-classified into unrelated bins.

**Figure 3:**
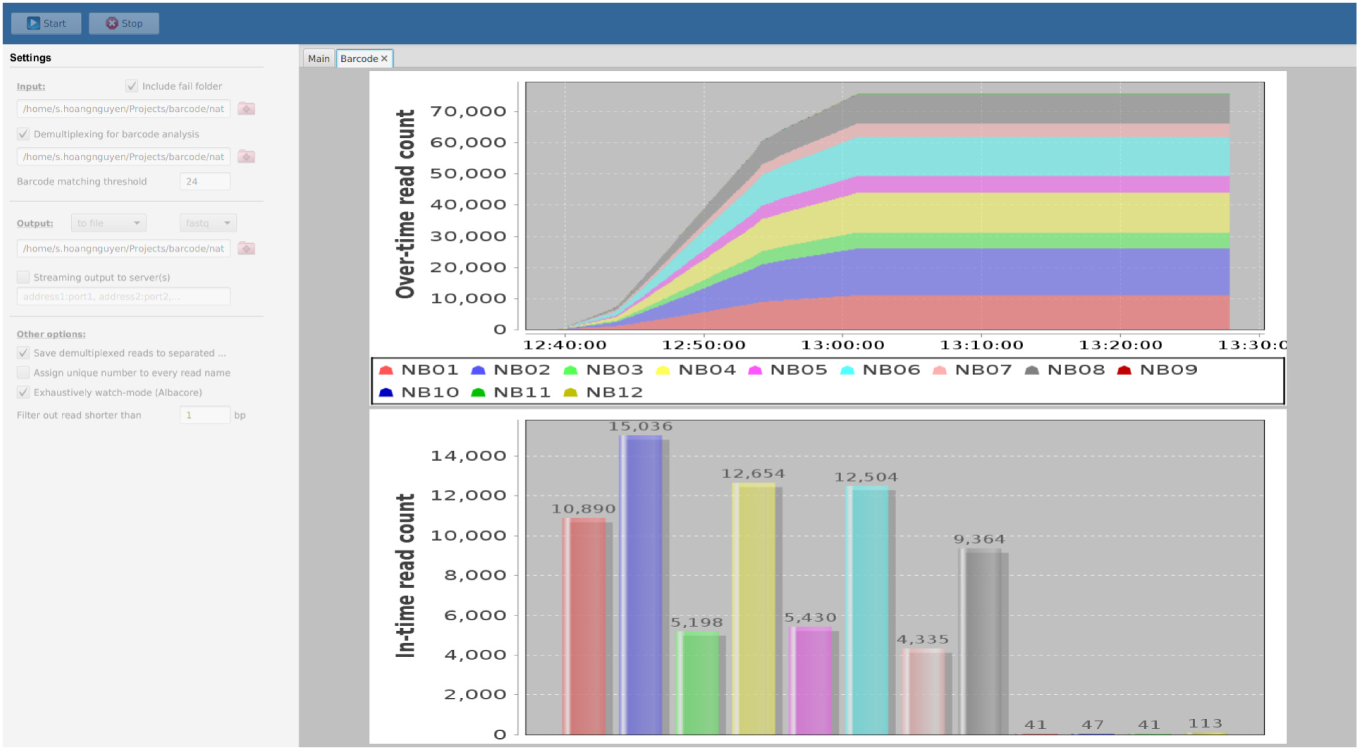
Graphical User Interface of npBarcode integrated in npReader. The result shown is for the a MinION run using Native barcoding kit on 8 libraries.

## 3. Conclusion

We have reported npBarcode, a tool supporting real-time demultiplex of nanopore sequencing data. Depending on requirements, users can choose to run the dedicated demultiplexer from command line or using it as part of npReader’s graphical user interface. The tool provides practitioners a flexible option to monitor a barcoded sequencing run as well as to integrate pooled sequencing into a streaming analysis pipeline.

